# Applying cis-regulatory codes to predict conserved and variable heat and cold stress response in maize

**DOI:** 10.1101/2021.01.15.426829

**Authors:** Peng Zhou, Tara A. Enders, Zachary A. Myers, Erika Magnusson, Peter A Crisp, Jaclyn Noshay, Fabio Gomez-Cano, Zhikai Liang, Erich Grotewold, Kathleen Greenham, Nathan Springer

## Abstract

Changes in gene expression are important for response to abiotic stress. Transcriptome profiling performed on maize inbred and hybrid genotypes subjected to heat or cold stress identifies many transcript abundance changes in response to these environmental conditions. Motifs that are enriched near differentially expressed genes were used to develop machine learning models to predict gene expression responses to heat or cold. The best performing models utilize the sequences both upstream and downstream of the transcription start site. Prediction accuracies could be improved using models developed for specific co-expression clusters compared to using all up- or down-regulated genes or by only using motifs within unmethylated regions. Comparisons of expression responses in multiple genotypes were used to identify genes with variable response and to identify *cis-* or *trans*-regulatory variation. Models trained on B73 data have lower performance when applied to Mo17 or W22, this could be improved by using models trained on data from all genotypes. However, the models have low accuracy for correctly predicting genes with variable responses to abiotic stress. This study provides insights into *cis*-regulatory motifs for heat- and cold-responsive gene expression and provides a framework for developing models to predict expression response to abiotic stress across multiple genotypes.

**One sentence summary:** Transcriptome profiling of maize inbred and hybrid seedlings subjected to heat or cold stress was used to identify key cis-regulatory elements and develop models to predict gene expression responses.

## Introduction

Plants are regularly exposed to variable environmental conditions throughout their life cycle and must be able to respond and acclimate to these conditions in order to survive and reproduce. Recent and rapid changes in climate have led to an increased frequency of extreme temperature fluctuations (Madani et al., 2018). Plants have developed sophisticated mechanisms at the cellular and metabolic levels that allow them to withstand temperature stress. In recent years, various regulatory mechanisms that involve hormone signaling, light signaling, circadian clock regulation and reactive oxygen species (ROS) homeostasis at transcriptional, epigenetic and post-translational levels during cold and heat stress have been identified (Chinnusamy et al., 2007; Nakashima et al., 2009; Mittler et al., 2012; Ohama et al., 2017; Li et al., 2018; Guo et al., 2018a; Ding et al., 2019, 2020).

Key players in the ability of plants to respond to temperature stress have been identified over the past decades. Members of CBF/DREB1 (C-repeat-binding factor/dehydration-responsive element-binding protein 1) TF family have been identified as essential regulators of the plant response to cold (Agarwal et al., 2006) by promoting cold stress responses through activating both cold-regulated (COR) genes and secondary signaling pathways (Fowler and Thomashow, 2002; Shi et al., 2017; Ding et al., 2019). Similarly, a family of HSFs (Heat Stress Transcription Factors) were identified as master regulators during heat stress by activating HSR (heat stress responsive) genes (Scharf et al., 2012) and turning on heat shock proteins (HSPs) that act as molecular chaperones, protecting cellular proteins by preventing their denaturation and aggregation, and facilitating the refolding of proteins damaged by heat (Wang et al., 2004; Busch et al., 2005; Charng et al., 2007). In addition, The AP2/EREBP (APETALA2 / Ethylene Responsive Element Binding Protein) gene family and its largest subfamily - the ethylene response factors (ERFs), participate in many developmental processes and play pivotal roles in adaptation to biotic or abiotic stresses including both cold and heat stress responses (Dietz et al., 2010; Mizoi et al., 2012; Hsieh et al., 2013; Cheng et al., 2013; Licausi et al., 2013; Yao et al., 2017; Huang et al.).

Functional binding of the TF requires both the presence of a matching motif (TF binding sites) and a proper chromatin state (Weirauch et al., 2014). Recent studies on accessible chromatin or chromatin modifications have documented many potential *cis-*regulatory elements in maize (Ricci et al., 2019; Oka et al., 2017); however, surveys of potential regulatory elements in unstressed plants have likely missed key potential stress-induced regulatory elements. The location of regulatory elements can be inferred through analyses of accessible chromatin, especially when focused on regions that are accessible only following stress treatments (Maher et al., 2018; Raxwal et al., 2020; Han et al., 2020). Alternatively, DNA methylation signatures are stable across tissues, developmental stages and environmental conditions, and can provide effective filters in mining functional regulatory elements (Crisp et al., 2020).

Several prior studies have documented gene expression changes in response to thermal stress in maize seedlings (Waters et al., 2017; Zhang et al., 2017; Li et al., 2017; Avila et al., 2018; Hoopes et al., 2019; He et al., 2019; Frey et al., 2020). These studies have found evidence for transcriptome changes in many of the expected pathways, and identified TFs that are up-regulated in response to heat or cold stress at multiple developmental stages. While great progress has been made in understanding some of the key TFs and TFBS motifs that play a role in response to heat and cold stress through studies of single varieties, the use of genetic variation within species can provide insights into the diversity of potential mechanisms by which variable *cis*-responses arise. Several studies have assessed variable responses to drought stress to reveal wide-spread *cis*-regulatory variation (Cubillos et al., 2014; Lovell et al., 2016; Liu et al., 2020). Analyses of maize allele-specific responses to several stress treatments also find evidence for both *cis*- and *trans*-regulatory variation (Waters et al., 2017). These studies have revealed that, although there are many genes with consistent responses to abiotic stress in multiple genotypes, there are also genes with highly variable responses to abiotic stress within the same species, and often this arises due to *cis*-regulatory variation. Promising results have been made to link *cis*-regulatory variation with gene expression and even plant phenotypes (Kwon et al., 2019; Alonge et al., 2020). However, it still remains largely unknown how sequence differences contribute to differences in responsiveness of promoters in general.

Machine learning approaches have provided new and powerful ways for understanding and predicting gene expression in plants (Azodi et al., 2020b; Wang et al., 2020b; Washburn et al., 2019). These approaches have been used to predict expression levels (Washburn et al., 2019; Sartor et al., 2019), regulatory architecture (Mejía-Guerra and Buckler, 2019), as well as gene expression responses to abiotic stress (Zou et al., 2011; Uygun et al., 2017, 2019; Azodi et al., 2020a; Schwarz et al., 2020). These studies highlight the potential to develop predictive models that use putative cis-regulatory motifs to predict gene expression responses to stress. However, whether these models can be applied to different genotypes other than the reference to predict consistent or variable expression response is not well understood.

We sought to investigate potential avenues for understanding transcriptome responses to heat and cold stress in several maize genotypes. Many genes that exhibited altered expression after a heat or cold stress event were identified and classified into clusters based on dynamic changes during a time-course sampling. These differentially expressed genes and clusters of genes with particular response patterns were mined to identify potential *cis*-regulatory motifs. These motifs were also used to develop machine learning models to predict responsiveness of gene expression. Our findings highlight parameters that are important in motif detection and modeling of expression responses. We found both potential uses, and pitfalls, in how data from one genotype can be used to predict expression responses in other genotypes.

## Results

### Dynamics of gene expression responses to heat and cold stress in seedling leaves

A series of experiments were conducted to provide insights into the *cis*-regulatory mechanisms that influence gene expression responses to abiotic stress in several maize genotypes. Maize seedlings were subjected to heat or cold stress experiments and RNA-Seq was performed at several time points during or after the stress (Figure S1). A time-course (TC) experiment sampled gene expression at nine time points in three maize genotypes that have *de novo* assembled genome sequences: B73, Mo17 and W22. Prior studies have shown differential responses of these genotypes to similar cold stress treatments (Enders et al., 2019; Waters et al., 2017). A second experiment (HY for hybrids) used three biological replicates to monitor gene expression at an early (1h) and later (25h) time point of the stress treatment that are both sampled at similar circadian timepoints. This HY experiment included the three inbred lines used for the TC experiment as well as F1 hybrids derived from crossing these parents. The third experiment (NM) used a single biological replicate to monitor gene expression at the early and late time points in a panel of 25 maize inbreds under control and cold stress conditions. In total, nearly 300 samples were generated and used to perform RNA-Seq (Table S1).

We initially focused on characterizing gene expression responses in the replicated HY experiment (Figure 1). A t-SNE analysis of the count per million (CPM) for all genes in all samples reveals a major genotypic effect as well as an effect of treatment/time point (Figure 1A). Control samples at 1h and 25 h cluster together as expected, both showing a slight shift from time 0 control samples. Cold samples at 1h are similar to the control 0h samples, both showing differences with control 1h samples which may reflect slowed circadian transitions in cold-stressed plants (Riva-Roveda et al., 2016). In contrast, heat treatment triggers a much faster response, as the heat 1h samples already show an obvious shift from control samples (Figure 1A). The 25h cold and heat stress treatments both are distinct from the control conditions. For each stress treatment, we identified differentially expressed genes (DEGs) by comparing the stress treated sample with a control sample isolated at the matched time point (“timeM_control”, Figure 1B). In addition, we also identified DEGs by comparing the stress treated sample to a control sample collected prior to the initiation of the stress treatment (“time0_control”, Figure 1B). It is worth noting that DEGs based on transcript abundance could reflect either differences in transcription or stability of transcripts and both mechanisms likely affect transcript abundance following abiotic stress treatments. We were interested in focusing on a set of genes that robustly respond to the cold or heat stress and for subsequent analyses will focus on genes that show consistent differential expression relative to both time 0 and the matched time-point (shown in red in Figure 1B).

**Figure 1.**
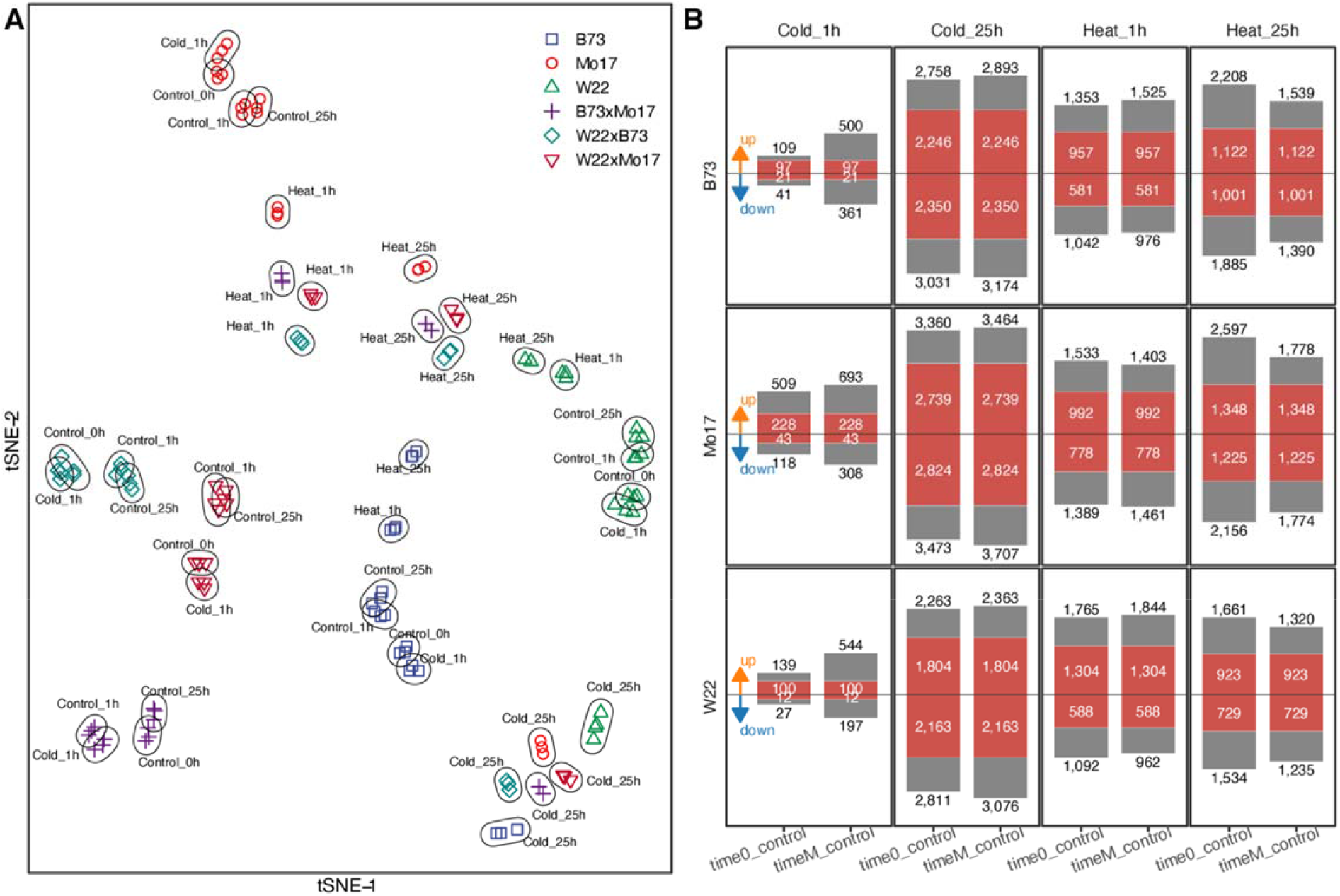
Identification of genes that are differentially expressed in response to heat or cold stress. (A) t-SNE clustering of samples from the hybrid (HY) experiment under control, cold and heat conditions. The genotypes are indicated by different symbols/colors and the conditions for each set of samples are indicated in the plot. (B) The number of differentially expressed genes (DEGs) under cold and heat conditions at 1 and 25 hour time points. For each time point the number of genes that showed differential expression (see Methods) relative to the control sampled at time 0 (the onset of the stress; time0_control) is shown as well as the number of genes that showed differential expression relative to the control sample collected at matching time point (i.e., 1h or 25h; timeM_control). Numbers inside red bars represent genes that are differentially expressed in both comparisons.

### Identification of patterns of gene expression response to heat/cold stress

The replicated data from the HY experiment was useful in identifying DEGs at early or later time points during a stress treatment, but the two sampled time points (1h and 25h) were insufficient to reconstruct the dynamics of transcript abundance changes upon stress treatment. To better understand the dynamics of stress response, we utilized the time-course sampling for the three inbred lines under stress treatments. Samples overall exhibited clustering that is consistent with genotype and type of treatment (Figure S2). Starting from the set of genes identified as differentially expressed at 1h or 25h under cold or heat conditions in at least one of the three genotypes, we further defined co-expression clusters based on their time-course expression profiles (see methods for details) (Figure S3). We focused on a subset of seven co-expression clusters which exhibit relatively stable expression in control samples but exhibit consistent patterns of up- or down-regulation under stress conditions with specific patterns of response (Figure 2, S3). In the response to cold stress, we identified four co-expression clusters that account for 39% of the DEGs and these represent groups of genes with early or late activation or repression in response to cold (Figure 2A). Three co-expression clusters were selected for response to heat stress accounting for 41% of the heat DEGs and these include transient activation, early activation and a set of genes with early repression (Figure 2A). The transient activation (0.5h) group includes genes that respond strongly in the first 30 minutes of the heat stress, but return to basal expression transcript levels at 1-2h of stress (Figure 2A), while the other heat activation group shows slightly slower activation and remains elevated for several hours (Figure 2A). These time-course co-expression clusters reveal notable differences in the response to heat and cold stress of maize seedlings. The response to heat is quite rapid and many genes return to basal levels while the response to cold stress is slower but often more persistent (Figure 2A and S3). This is consistent with the observation that heat 1h samples shifted away from control much faster than the cold 1h samples (Figure 1A). A GO analysis performed using all up-regulated genes or the up-regulated genes for B73 within each cluster reveals enrichment for terms associated with transcription regulation for the cold response and response to heat or protein folding for heat stress response (Figure 2B).

**Figure 2.**
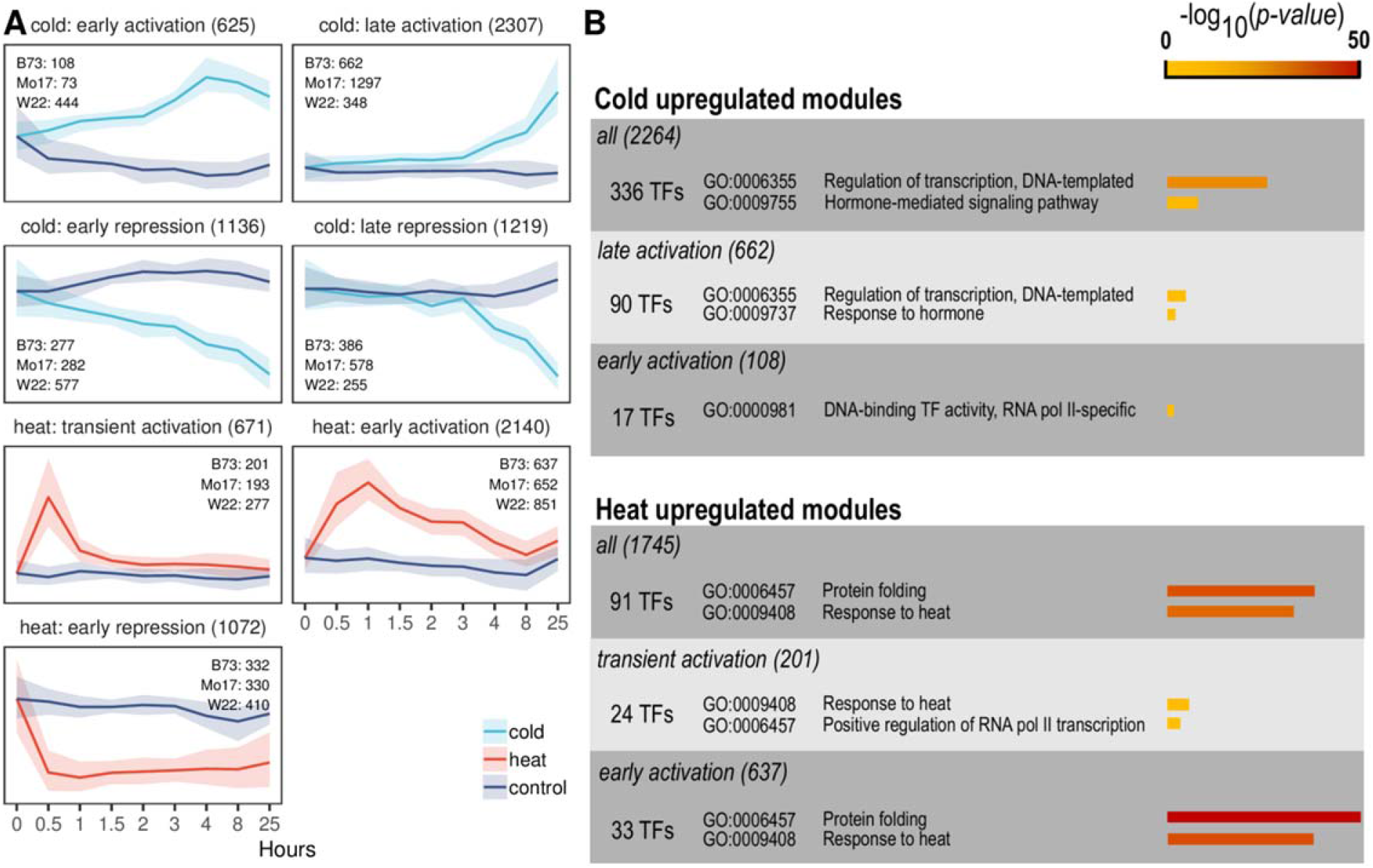
Expression profiles and GO enrichment of cold- and heat-responsive gene clusters. B73, Mo17 and W22 genes that exhibit significant DE after 1 hour or 25 hours stress treatment were used to perform co-expression clustering based on their time-course expression pattern (see methods). (A) The expression pattern for the most abundant co-expression clusters that represent early, late or transient changes upon cold or heat treatment. The median expression level of control and stress conditions for the genes within each module is shown and the number of B73, Mo17 and W22 genes in each module is indicated. Ranges at each time point represent 25%-75% quantile expression levels. (B) For the set of all up-regulated genes under cold or heat conditions we determined the number of TFs and assessed GO enrichment. Two highly enriched GO terms (with significance p < 0.01) are shown for each module, and the significance level is indicated.

Given the importance of transcriptional regulation in the control of stress responses (Ohama et al., 2017), we identified TFs that are up- or down-regulated in response to heat or cold (Figure 2B). The analysis of all up-regulated TFs or the subsets found in specific co-expression clusters revealed that some TF families exhibit enrichment for responses to these stimuli (Figure S4). Eleven TF families are enriched in the set of all cold up-regulated TFs including EREB, WRKY and NAC families (Figure S4). While each of these families are enriched in the full set of up-regulated genes, some exhibit stronger enrichment in the early or late activation clusters. Fewer (six) families of TFs are enriched in genes up-regulated in response to heat with the HSF family being the most significant (Figure S4B). There are 29 HSF TFs in the B73 genome (Yilmaz et al., 2009; Zhang et al., 2020) and 10 of these are up-regulated in response to heat (Figure S5A-B). Many of the HSFs that are classified as DEGs exhibit very rapid increases in transcript levels (some with >100-fold increases in 30 minutes) (Figure S5). We also noticed that several of the HSFs (*ZmHSF4, ZmHSF12, ZmHSF20*) that are not classified as DEGs actually exhibit very strong activation at 30 minutes, but have already gone back to basal expression levels by 1h, and are not classified as differentially expressed in our replicated experiment (Figure S5D). We also assessed the expression patterns for a set of 104 TFs that have been previously reported to play a role in cold stress response in various cereal species (Baillo et al., 2019) and found that 40 of these TFs are up-regulated in our B73 dataset and include examples in many different co-expression clusters (Figure S6).

### Identification of enriched motifs for each cluster of heat/cold stress induced genes

We sought to identify DNA motifs associated with responsiveness to heat or cold stress by assessing sequences of all DEGs or specific co-expression clusters for significantly enriched motifs using STREME (Bailey, 2020) (Figure 3). Determining the ideal search space of potential regulatory elements when searching for motifs associated with transcript abundance responses can be challenging. While many TFBSs are likely quite near the transcription start site (TSS), they could also be located in more distal regions. The identification of enriched motifs within B73 (B) or B73/Mo17/W22 (BMW) DEGs was performed using several sets of search spaces including regions of different promoter size (0.5kb, 1kb and 2kb), orientations (upstream of TTS, downstream of TSS or a combination of both), and a subset of the sequence filtered to only retain regions classified as unmethylated (Crisp et al., 2020) or accessible chromatin [based on (Ricci et al., 2019)] (Figure 3A, see methods for details). A total of 1,950 motifs were identified using B73 DEGs and 4,981 unique motifs were obtained using BMW DEGs with the number of motifs varying substantially for different groups of DEGs (Figure 3B; Figure S7). A similar analysis was also performed using the DREME algorithm (Bailey, 2011). Substantially more motifs were found using DREME, but these often included short motifs and provided lower prediction accuracies in subsequent analyses so we focused on the motifs identified using STREME.

**Figure 3.**
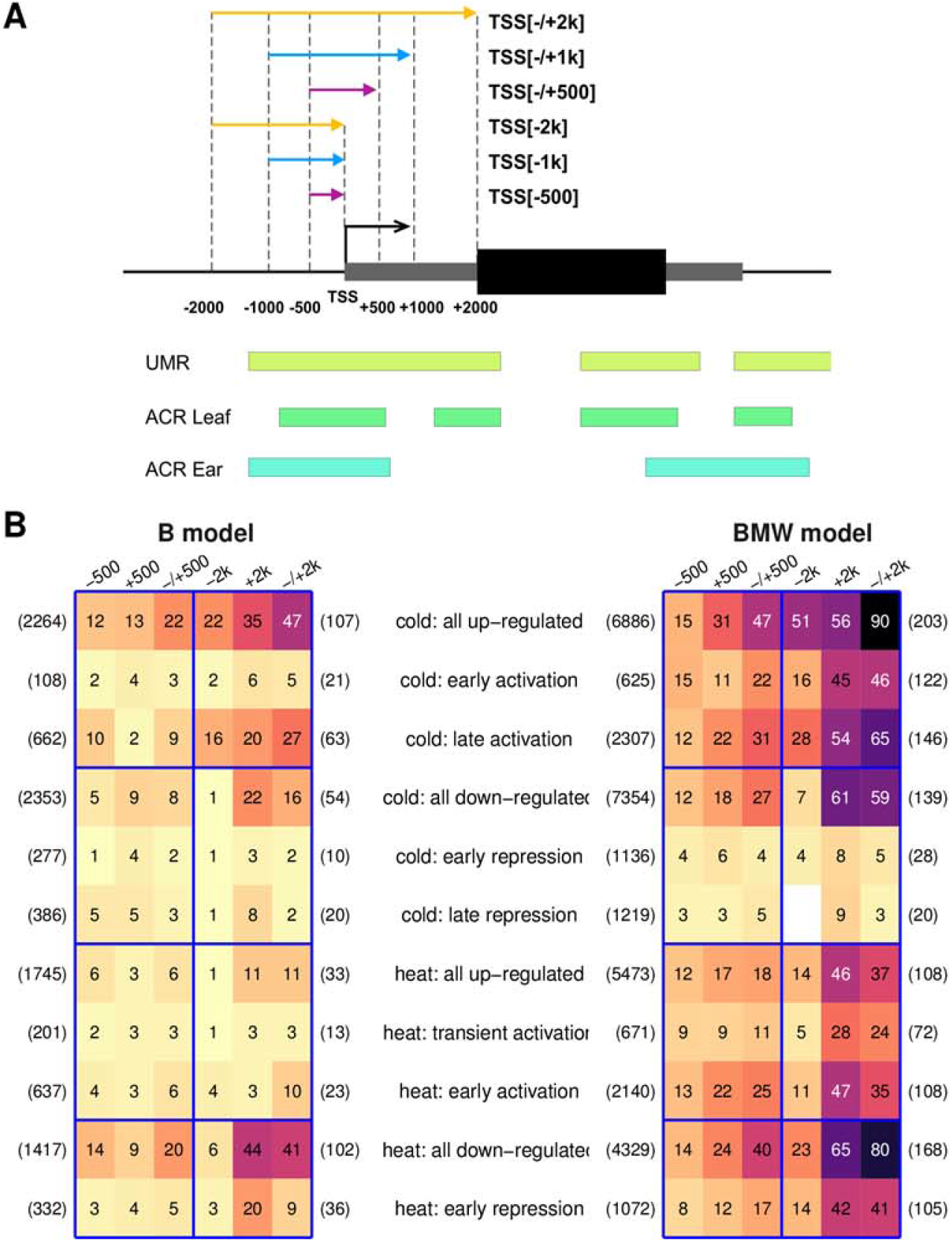
Identification of enriched motifs in cold- and heat-responsive genes. (A) Varying potential ‘promoter’ sequence spaces were utilized to search for motifs that are enriched in different sets of genes. The schematic indicates a representative gene with the transcription start site (TSS) indicated. The potential regions include different lengths of sequences upstream the promoter (−500, −1k, −2k) as well as sets of sequence that include both upstream and downstream sequence (i.e., +/−500). In addition, for each of these potential regions we also subset the sequence to only include regions that are unmethylated (UMR - unmethylated regions) or that are classified as accessible based on ATAC-seq analysis (ACR - accessible chromatin region) in leaf or ear tissue (Ricci et al., 2019). (B) The number of non-redundant motifs found using B73 promoter spaces (“B model”) or B73/Mo17/W22 promoter spaces (“BMW model”) and different search spaces is indicated. Colors indicate numbers of motifs identified (the darker, the more motifs found) and correspond to numbers in each cell. Numbers in parentheses on the left side of the heatmap indicate the number of genes used for motif mining; while numbers on the right side of the plot indicate the total number of non-redundant motifs found for each set of genes. More detailed search results using different promoter sizes along with different chromatin properties (UMRs, ACRs) are shown in Figure S7.

As expected, we tended to find more motifs when assessing large groups of genes or using larger sequence space (Figure 3B, S7). In many cases we identified similar numbers or potentially higher numbers of motifs when using the sequences downstream of the TSS compared to solely using upstream sequences (Figure 3B). This may reflect transcriptional regulatory elements that are present within transcribed sequences (5’UTRs, introns, exons) or the presence of motifs within RNA that affect stability of transcripts. When we restricted the search space to only include the unmethylated or accessible regions, we found substantially fewer enriched motifs (Figure S7). Although fewer motifs were identified in these restricted regions, we found that these often were the same motifs identified in the full sequence and often had higher fold-enrichment.

The full set of 1,950 B73 motifs were collapsed into 558 motif groups based on sequence similarity, 143 of which can be assigned to known transcription factor binding sites (TFBSs) in cis-BP (Weirauch et al., 2014), while the remaining 415 are novel. An assessment of the top 40 most significantly enriched motifs from the search that used +/−2kb sequences for each co-expression cluster reveals that a subset of the enriched motifs reflect previously characterized TFBSs while others represent novel motifs not captured by the current collection of plant TFBSs (Figure S8). The motifs that represent known TFBSs include some of the expected sites such as HSF binding sites for heat or CBF/DREB binding sites for cold (Figure S8). Analysis of the motifs that are identified using all three genotypes finds many of the same motifs identified using B73 only (Figure S8). The analysis of the enrichment of several motifs for each of the clusters reveals examples in which some motifs are more strongly enriched in co-expression clusters and other examples that exhibit similar enrichment for the co-expression clusters and all up- or down- regulation genes (Figure S9). A primary goal for the identification of motifs was to collect a set of features that may be useful to develop models to predict response to heat or cold in maize. In this application it is not critical that all motifs be valid as the models can be trained to utilize the most predictive features.

### Generation of models to predict heat- and cold-responsive gene expression

We sought to generate and assess predictive models of B73 stress-responsive expression using the potential *cis*-regulatory elements (motifs) as “features” to classify a gene’s expression response to heat or cold stress. The full set (all motifs associated with the sets of all up- or down- regulated genes or specific co-expression clusters) of motifs identified for cold or heat responsive expression (above) were used and we determined the presence/absence for each motif within different sets of search spaces (Figure 3A) to assess how the use of different potential ‘promoter’ regions would affect model performance. The approach previously described by Zou et al (Zou et al., 2011; Uygun et al., 2017, 2019; Azodi et al., 2020a) was implemented to utilize the motif features to predict whether genes will exhibit cold- or heat-responsive expression. For each co-expression cluster, we compared the genes in the cluster to an equivalent number of genes that are expressed, but not classified as DE; and ran the model 100 times using all motifs, or only retaining the most highly enriched motifs (top 30, 50, 100 or 200) followed by calculation of the AUROC (see Methods for details; Figure 4; Figure S10).

**Figure 4.**
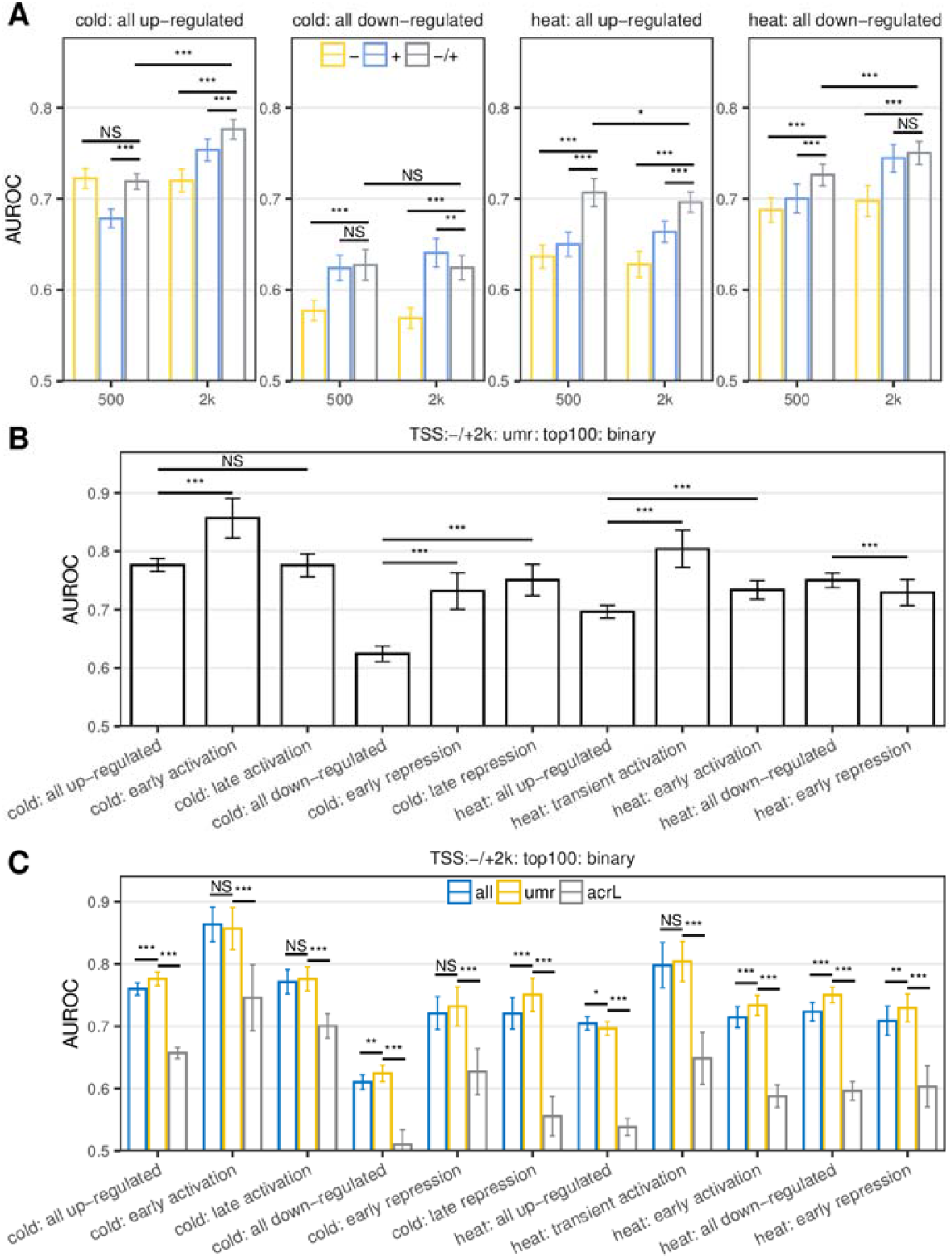
Performance (Area Under ROC Curve, AUROC) comparison of different machine learning models predicting cold and heat responsive expression. (A) The performance comparison of models trained using small promoter spaces (−500, +500, −/+500) against larger promoter spaces (−2k, +2k, −/+2k). In each case the same chromatin filters (UMR) and number of features (top100) were used for training. (B) The performance comparison of models trained using the full set of DEGs against models trained using specific co-expression clusters. In each case same sized promoter spaces (−/+2k), chromatin filters (UMR) and number of features (top100) were used. (C) The performance comparison of models using all genomic sequence (“all”), UMR regions only (“umr”) or using leaf accessible regions only (“acrL”) with the same sized promoter spaces (−/+2k) and number of features (top100). In each training average AUROC (N=100 model runs) is shown along with the standard deviation. Pairwise comparisons were made using t-test with significance levels indicated (*: 0.05; **: 0.01; ***: 0.001)

Overall, we were able to achieve moderate accuracies for predictions of which genes would respond to heat or cold stress in B73 with higher accuracies for cold up-regulated genes and heat down-regulated genes (Figure 4; Figure S10). In general, the use of sequences both upstream and downstream of the TSS (+/−2kb, +/−500bp) provides higher prediction accuracies than using the upstream or downstream sequences alone (−2kb or +2kb, −500bp or +500bp), and using longer sequences surrounding the TSS (+/−2kb) provides higher prediction accuracies than the use of only +/−500bp sequence (Figure 4A, S10A). For subsequent analyses we assessed several additional parameters while using motifs within 2kb of the TSS. A comparison of the prediction accuracy for models developed for specific co-expression clusters compared to models developed for all up- or down-regulated genes reveals significantly improved performance for genes that are down-regulated in response to cold, early-activation in response to cold or transient activation in response to heat (Figure 4B). However, there was no significant increase for groups of genes with late activation in response to cold or down-regulated in response to heat (Figure 4B). We tested whether only using motifs present within unmethylated regions or accessible chromatin could improve prediction accuracy relative to using all sequences within the +/−2kb window (Figure 4C). The prediction accuracies increased significantly for 6 of the 11 groups tested when focused only on the motifs within UMRs, even though substantially less sequence was used (Figure 4C). In contrast, the use of motifs only with accessible regions, based on a prior study of maize seedling tissue (Ricci et al., 2019), exhibits significantly decreased performance for nearly all of the groups of genes (Figure 4C). A comparison of the performance of models that simply utilized presence/absence of the motif (“binary encoding”) compared to models that included the number of motifs per gene (“quantitative encoding”) did not find major differences, although the binary model often has slightly improved performance (Figure S10B). Finally, we also assessed the relative performance of models that utilized only the top 30, 50, 100 or 200 motifs for each group of genes and did not find substantial improvement when using higher numbers of motifs suggesting the top 30 motifs often capture most of the critical information (Figure S10C). For many of the groups of genes using only the top 30 or 50 motifs provided similar performance compared to larger sets of motifs (Figure 4D).

The motif enrichment scores were compared with the feature importance of that motif in the predictive models for each of the sets of genes that respond to heat or cold stress (Figure S11). We were interested in determining whether the most highly enriched motifs were also the most predictive features. For the genes that are up-regulated in response to heat the HSFC1 motif was the most enriched and was also the highest feature importance score in models (Figure S11). In contrast, for the other groups of genes (down-regulated in response to heat and up / down regulated in response to cold) the motif enrichment score was not well correlated with the feature importance score (Figure S11). Often, the motifs with the highest feature importance scores had moderate enrichments ranks.

### Genetic variation for responses to cold and heat stress

In the analyses above we focused on generation of predictive models based on the B73 genomic sequence and changes in expression in B73. However, a key goal is to better understand and predict variable responses to abiotic stress in different genotypes. Our experimental design allowed us to characterize the variation in cold or heat responsive expression in three maize inbreds and to separate *cis-* and *trans*-acting regulatory variation using the F1 hybrids. Genes that are differentially expressed (relative to both time 0 and a matched time point) at 1hr or 25hr of heat or cold stress in at least one of the three inbred genotypes were used to classify variable responses. The results for 25h cold up-regulation are shown in Figure 5 while the full sets are shown in Figures S11-13. A comparison of the genes identified in each genotype revealed that while there is a subset of genes that exhibit consistent up-regulation in all three genotypes there are also many examples showing up-regulation in only one genotype (Figure 5A; S12A). Hierarchical clustering of the genes that only exhibit significant up-regulation in one or two of the genotypes revealed that many of these show minimal expression changes in some genotypes (Figure S12B).

**Figure 5.**
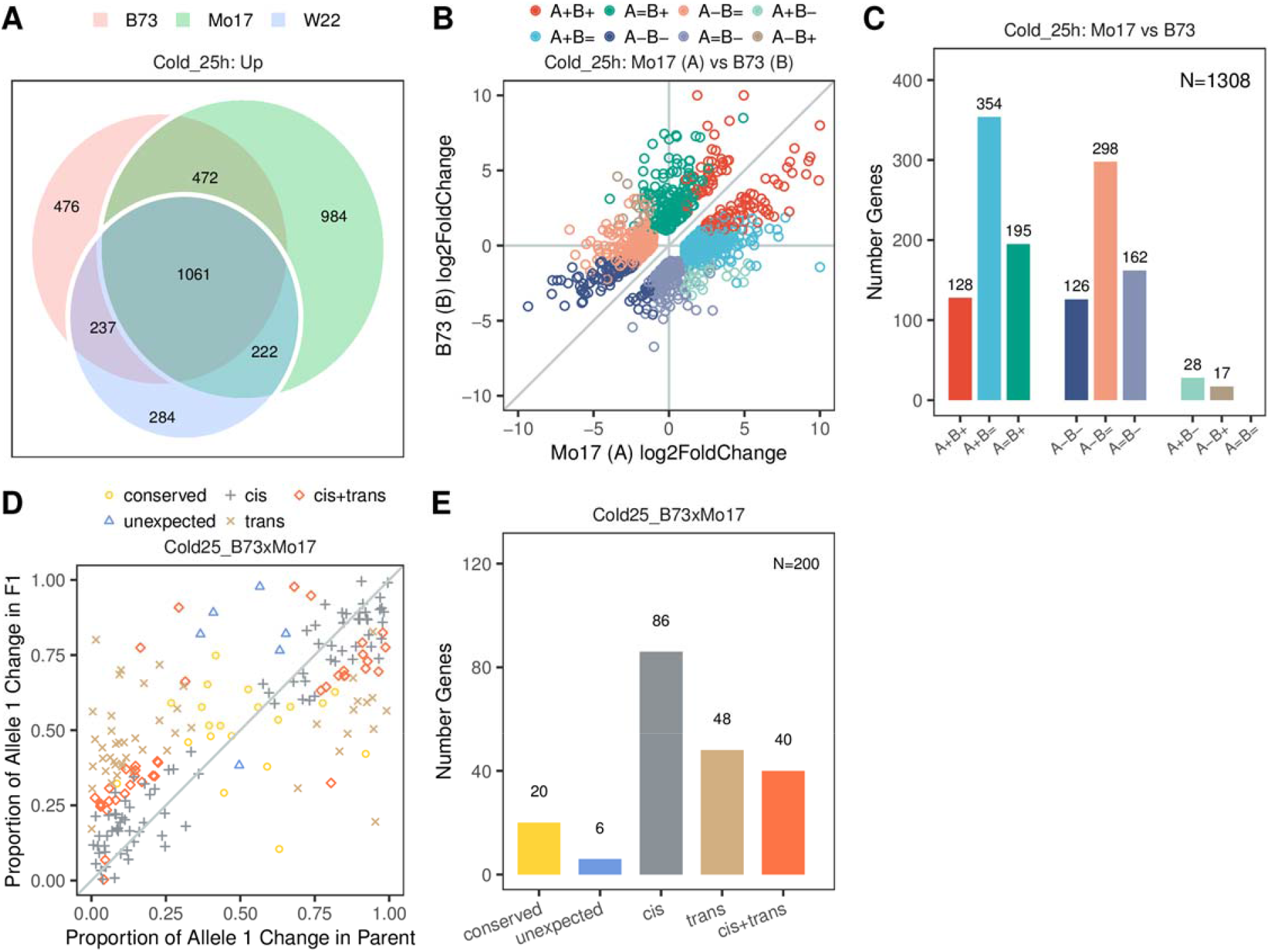
Characterization of genes with variable stress responsive patterns among inbreds. (A) A Venn diagram is used to compare the genes that are up-regulated in response to 25h of cold stress for B73, Mo17 and W22. The overlap of DE genes at other time points is shown in Figure S11A. (B) For genes that show significantly stronger (or weaker) response to cold at 25h in B73 compared to Mo17 we show the log2 fold-change (cold 25h / control 25h) for both inbreds. The classification of differential responses for other genotype contrasts and timepoints is provided in Figure S12. Post-hoc tests were used to classify (colors) genes with varying differential expression between genotypes; red indicates up-regulation in both inbreds (with different response levels), green indicates genes only up-regulated in Mo17, etc. The number of genes in each category is shown in (C). For each class the response in the two genotypes (A and B) is indicated as up-regulated (“+”), down-regulated (“-”) or not DE (=). (D) For the subset of genes classified as having a response in only one of the two genotypes that also had SNPs we assessed allele-specific expression in the F1 hybrid. The proportion of allele 1 (B73) change in stress vs control of the F1 (x-axis) was compared to the proportion of the change in expression in the parental genotypes (y-axis). A maximum likelihood model was applied to classify *cis-* and *trans-* inheritance patterns and these classifications are shown in different colors. The number of genes classified into each type of regulatory pattern for response to abiotic stress are shown in (E). Similar analyses for other genotypes, stress and time points are shown in Figure S13.

Our goal was to identify a set of genes that have a robust response to stress in some genotypes but not in others to identify true variable responses. We assessed the set of genes that exhibit a significant response in only one or two genotypes and introduced an interaction term (genotype:condition) to model the genotype-specific condition effect under the generalized linear framework of DESeq2 (Figure 5B-C, S13). This allowed us to classify genes that exhibit significant up-regulation in one genotype but not the other (A+B= or A=B+), as well as a set of genes that respond more strongly in one genotype but do exhibit responses in both genotypes (A+B+) (Figure 5B-C). We proceeded to assess allele-specific expression for the sets of genes that have a strong expression response in one genotype but not in another. A subset (11-37%) of these genes with response in one genotype but not another contain SNPs and sufficient allele-specific read-depth to perform allele-specific expression analysis in the F1 hybrid (Figure 5D-E; Figure S14). Using a model that incorporates allele-specific expression in the F1 and relative expression levels in the two parents (see methods for details), we determined their regulatory patterns as *cis-*only, *trans-*only, or a mix of *cis-* and *trans-* (Figure 5D-E; Figure S14). These analyses allow us to discover many examples of genes with variable response to heat or cold stress in these three genotypes. The allele-specific data revealed many cases of both *cis*- and *trans*-regulatory variation that controlled the differential response to stress. Interestingly, many of the differences reflect cis-regulatory variation, which suggests that changes in the promoter sequences for one allele alter the ability to respond to abiotic stress. It is worth noting that the number of genes identified with *cis*-regulatory variation is likely an under-estimate since many genes with differential response did not have SNPs within the coding region which precludes the ability to perform allele-specific expression analyses.

### Association of variable response to heat or cold stress with model predictions or features

We sought to assess how several factors might influence predictions of response to heat or cold stress across multiple genotypes (Figure 6). The initial models described above were trained using B73 DEGs and motifs. When these models were applied to DEGs and promoter sequence from Mo17 or W22 we found lower performance (Figure 6A). However, even in these cases the AUROC scores are greater than 0.7 for most of the groups of DEGs in the other genotypes suggesting that information from B73 based models can be useful for predicting responses in other genotypes. We then trained models that utilized DEGs from all three genotypes (as well as control genes from each genotype) and assessed the accuracies for prediction of DEs in each genotype. This approach provides increased prediction accuracies for all three genotypes with some groups of genes exhibiting AUROCs >0.95 (Figure 6A). While these accuracies were quite impressive, they likely reflect over-fitting due to the inclusion of common genes that are up-regulated in multiple genotypes. We further tested the ability for these models to correctly predict variation for responses to heat or cold stress. We used the classification of the relative responses in multiple genotypes (Figure 5C, S13) and assessed the prediction for these genes relative to the observed patterns (Figure 6B). The majority of genes that exhibit transcript abundance response in both genotypes are correctly predicted to respond in both genotypes and relatively few of these genes are predicted to not respond in either genotype. The proportion of these genes with conserved response that is correctly predicted is higher for the model trained on all three genotypes (89%) than for the model trained using only B73 data (69%). The correct prediction accuracies are much lower when focusing on the subset of genes that only respond in B73 or in another of the genotypes. For the genes that respond in B73 but not in the other genotype being compared, the model trained on B73 data correctly predicts 16% of these genes. Application of this same model to predict genes that would respond in another genotype but not in B73 results in lower rates of correct predictions (7%). The model trained using data for all three genotypes shows less difference in correctly predicting B73-specific responses (15%) compared to responses in the other genotype (11%), as expected (Figure 6B).

**Figure 6.**
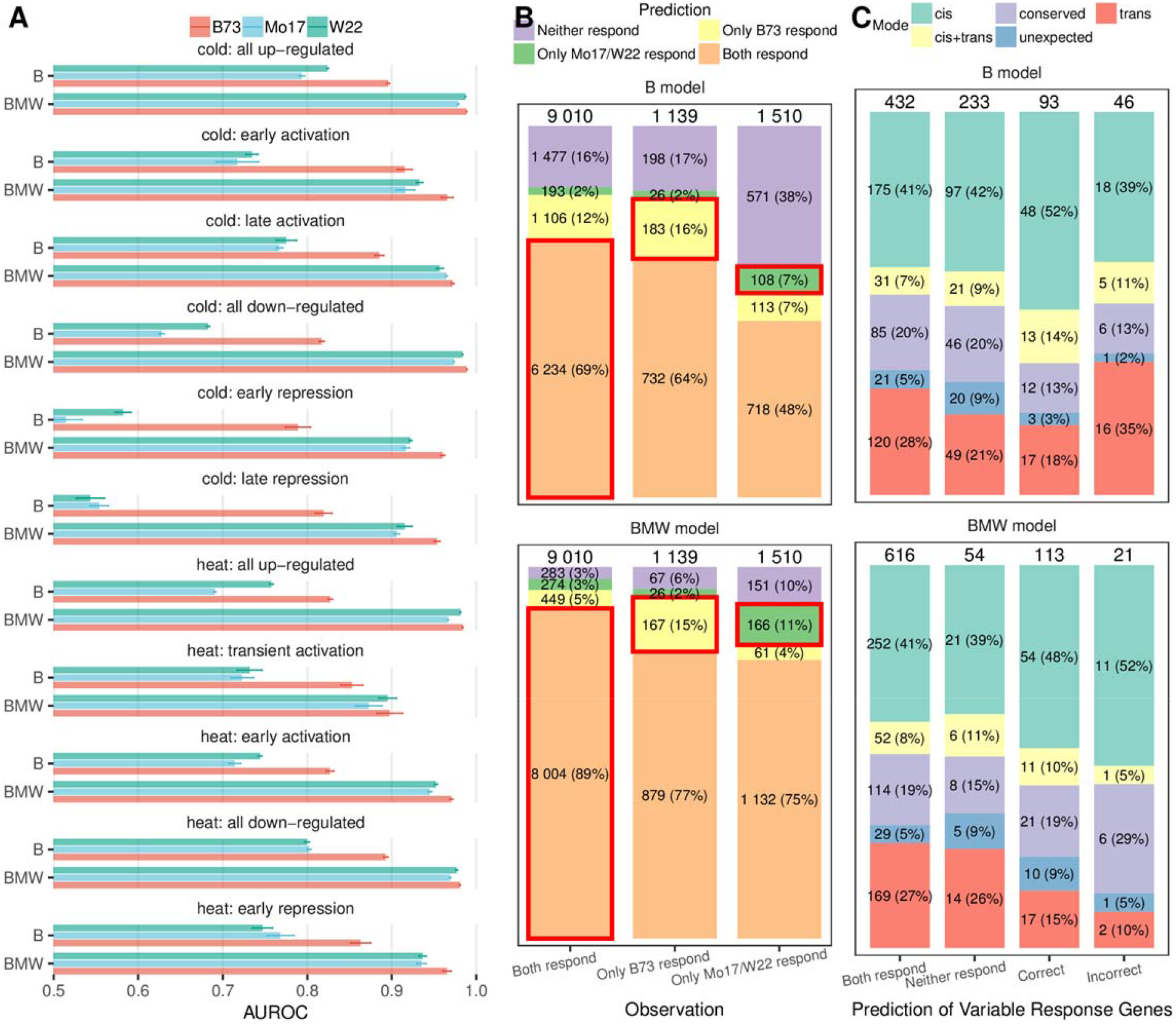
Cross-genotype performance of machine learning models predicting cold or heat responsive expression. Models were trained only using B73 sequence and DE labels (“B model”) or data from all three genotypes (“BMW model”). (A) AUROC for models predicting stress responsive expression in B73, Mo17 and W22. Average AUROC (N=100 model permutations) is shown along with the standard deviation for both B and BMW models. (B) Model prediction accuracy for genes showing consistent (“Both respond”) or variable (“Only B73 or Mo17/W22 respond”) response patterns among genotypes. In each observed category the number and proportion of predictions were marked in the plot with the correct predictions highlighted with red boxes. (C) Dissection of regulatory patterns for genes showing variable response patterns among genotypes. Variable response genes were first grouped by whether model prediction agrees with observed status (“Correct” if the model correctly predicts one genotype responds but the other does not, “Incorrect” if the model predicts oppositely, “Both respond” and “Neither respond” if the model predicts both or neither genotypes respond - although in reality only one genotype responds). Then within each group the number of proportion of different regulatory patterns (“*cis*”, “*trans*”, etc) were marked.

Regardless of which model is used, we find that most genes with genotype-specific responses were incorrectly predicted to respond in both genotypes (Figure 6B). Since our predictions are largely based on the presence of putative *cis*-regulatory motifs it would be expected that prediction accuracies should be higher for genes with *cis*-regulatory variation for responsiveness when compared with genes that exhibit *trans*-regulatory variation. The classification of *cis*- and *trans*- regulatory variations were assessed for genes with variable prediction in each of the groups (Figure 6C). As expected, genes with the correct predictions do show slightly higher proportions of *cis*-regulation and are depleted for *trans*-regulation. However, the enrichments are fairly subtle and many genes with *cis*-regulatory variation are still predicted incorrectly.

### Potential examples of TFBS variation associated with differences in cold-responsive gene expression

We collected expression data for a single replicate of control and cold stressed plants at two time points (1h and 25h) for 23 maize genotypes that all have genomic resources including SNP calls relative to B73. We used the expression and genomic data for these genotypes to associate local haplotype variation with expression changes for a subset of 2,147 genes (Figure 7). These include 1,088 up-regulated and 1,059 down-regulated genes that exhibit variable response in B73, Mo17 and W22 at 25h after cold treatment (Figure 5A-C). B73, Mo17 and W22 were included in this panel and exhibit expression changes that support the classification of variable responses (Figure S15A). In order to identify examples of clear classification as “response” or “no response” to cold we assessed the distribution of the ratio (log2) of transcript abundance in cold compared to control to identify genes with clear bi-modal distributions (Figure 7A-B). While many genes had unimodal distributions, there are a subset of 529 genes with significant bi-modal or multimodal distributions for which genes can be classified as responding to cold or not responding (Figure 7A-B). As expected based on the selection criteria of these genes, at least one of the three core genotypes (B73, Mo17 and W22) is always within both the responding and non-responding group.

**Figure 7.**
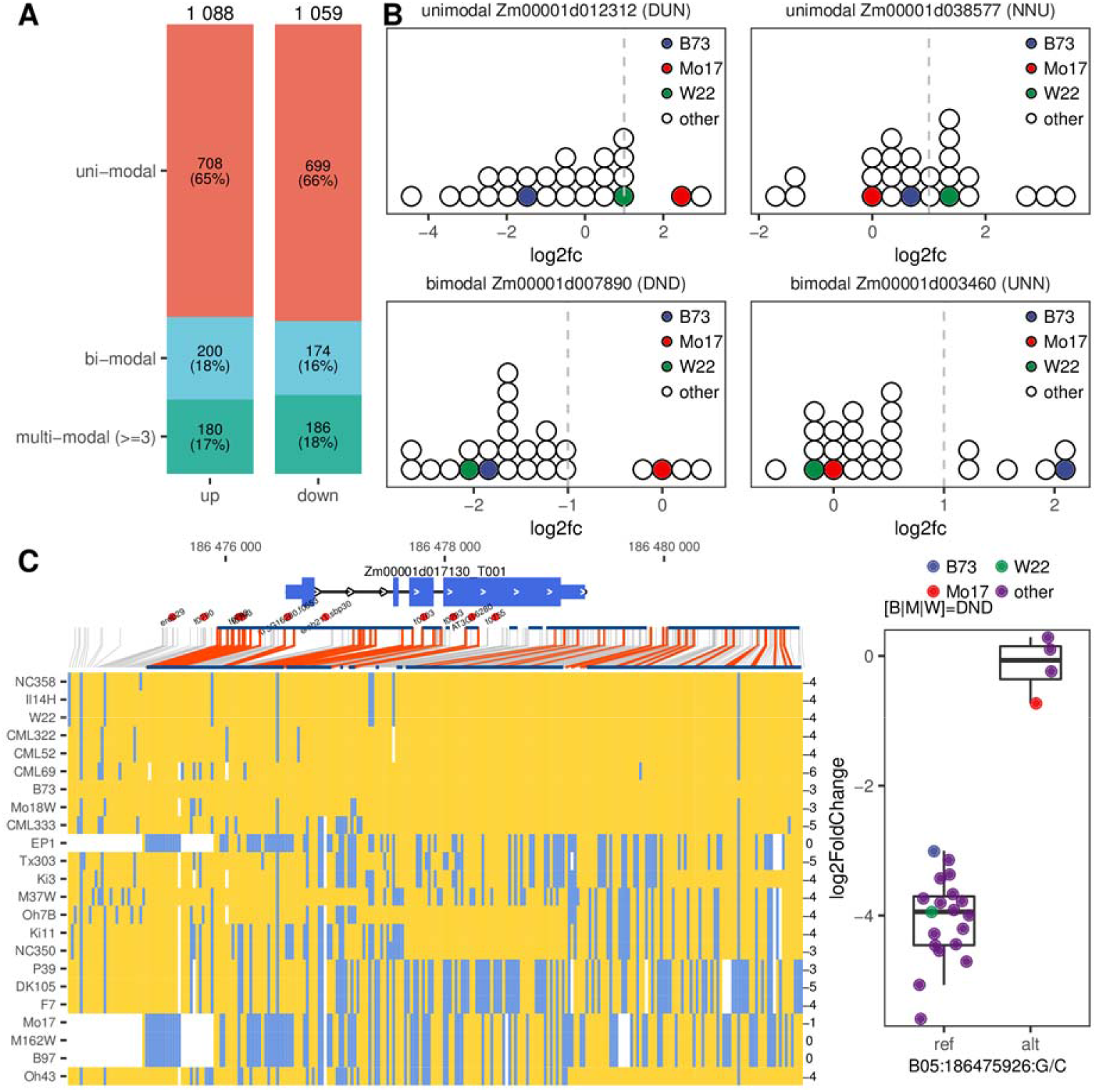
Identification of *cis*-regulatory variants associated with variable cold responsive pattern in a panel of 25 maize genotypes. (A) The proportion of ~2000 variable response genes showing uni-modal, bi-modal and multi-modal distributions of the log2 fold change ratios (cold/control); (B) Examples of two genes showing uni-modal log2fc ratios and two showing bi-modal ratio distributions; (C) *Cis*-regulatory variation associated with cold responsiveness in a maize mitochondrial transcription termination factor (mTERF17). Heatmap shows biallelic variants (SNP and short indels) within 2kb of the gene with yellow indicating reference (B73) allele and blue indicating alternate allele. Log2 fold change of each genotype is shown on the right of the heatmap. Below the gene structure plot are locations of the most (top30) enriched cold-responsive motifs (red dots) with the motif name marked, as well as haplotype blocks (dark blue segments) identified using PLINK. The boxplot on the right shows the top associating variant and the log2fc distributions of genotypes carrying the two alleles.

We then aligned the genomic sequences of the 25 maize genotypes to B73 and extracted bi-allelic variants within 2kb of each gene to perform a local association test with the cold response pattern (i.e., responding and non-responding genotypes). PLINK (Purcell et al., 2007) was used to perform a standard chi-square association test for each gene. Many of the genes (308/529) have at least one significant association of a local variant and the expression response. As expected, the majority (76%) of genes that are classified as having *cis*-regulatory variation exhibit a significant association with the local haplotype (Figure S15B). In contrast, a smaller proportion (50%) of genes classified as having *trans*-regulatory variation are significantly associated with local haplotype (Figure S15B). Several examples were assessed to visualize the haplotype variation relative to expression response and motifs (Figure 7C, S16). One interesting example is a maize mitochondrial transcription termination factor (mTERF17, Figure 7C), which showed significant down-regulation upon cold stress in all maize genotypes except Mo17, B97, EP1 and M162W. Studies in Arabidopsis suggested an important role of mTERF family members in the regulation of mtDNA transcription and disruption of some family members results in reduced mtRNA accumulation and altered organelle biogenesis (Kleine, 2012; Shevtsov et al., 2018). Local association tests identified a number of significantly associated variants including some in the promoter region overlapping the identified cold-responsive motifs (Figure 7C). It is likely that by accumulating these mutations the Mo17 haplotype/allele has lost the ability to respond to cold stress and thus mtRNA accumulation will be unaffected by cold stress in these genotypes. Other examples include a respiratory burst oxidase (rboh4) showing significant up-regulation in most maize genotypes except B73, CML333, NC350 and M37W, as well as a dihydroflavonoid reductase (dfr1) showing significant up-regulation in most maize genotypes but not in Mo17 and four other genotypes (Figure S16A-B). Local association tests also identified a number of potential cis-regulatory variants overlapping some of the most enriched cold-responsive motifs (Figure S16).

## Discussion

Understanding the gene expression responses to abiotic stress will contribute to efforts to develop more resilient crop varieties. In this study, we focus on monitoring transcriptome responses to heat and cold stress in maize seedling leaf tissue. The use of paired heat and cold stress with the same sampling times and genotypes revealed some key differences in the dynamics of response. Many of the responses to the heat stress event used here occur very rapidly and are fairly transient. In contrast, the cold stress employed here tends to have less drastic effects in the first hours, but includes changes in expression that continue, and often strengthen, over the course of 24h. This may reflect differences in how plants sense and respond to cold and heat stress and provides an opportunity to use quite distinct stress responses for attempting to understand important *cis*-regulatory features of response to abiotic stress.

A key long-term is to understand how *cis*-regulatory elements, and variation among them, drives variation for gene expression responses to thermal stress in different genotypes of crop species. This has implications for understanding the evolutionary sources of regulatory variation for stress responsiveness and for the ability to identify genotypes with particular responses to abiotic stress. The analyses performed in this study provide insights into the transcript abundance changes that occur in response to heat and cold stress, the applications of predictive models of gene expression responses as well as the potential opportunities and associated risks when attempting to predict expression responses across genotypes.

### Identification of TFs and TFBSs associated with response to heat and cold stress

Environmental stresses induce significant differential gene regulation, and these large-scale changes are likely accomplished through activation of a relatively small set of stress-responsive TFs that activate, or repress, a much larger number of stress-responsive genes. We identified a number of TFs that exhibit increased transcript abundance following heat or cold stress including a subset that are activated very early upon exposure to stress (Figure 2B, S5-6). In many cases we also observed connections between the TFs that were up-regulated and some of the TFBSs that were enriched in the promoters of up-regulated genes. In heat up-regulated genes, there was significant enrichment for both HSF TFs and a previously identified HSFC1 binding site (Franco-Zorrilla et al., 2014) in the early activation cluster, as well as the full set of heat up-regulated DEGs. In cold up-regulated genes, there was significant enrichment for both WRKY TFs and several previously identified WRKY binding sites, as well as a significant enrichment for both MYB TFs and several previously identified MYB binding sites (Franco-Zorrilla et al., 2014). Members of each of these TF families have demonstrated roles in stress responses. HSF TFs are thoroughly characterized regulators of heat stress responses (Scharf et al., 2012), while members of both WRKY and MYB TF families have been implicated in cold stress responses (Chen et al., 2012; Li et al., 2015). Finally, the identified TF family enrichments (Figure S4) appear to nearly completely mirror previous observations of cold and heat responsive TF families (Li et al., 2019, 2020), though several additional family enrichments were discovered in our module-based analyses.

### “Promoter” definitions influence ability to identify motifs and predict responses

Our understanding of the architecture of regulatory regions in plants remains limited (Weber et al., 2016; Long et al., 2016). It is tempting to use a simple criterion such as the 1kb of sequence immediately upstream of the core promoter when searching for potential regulatory elements. However, there is evidence that important regulatory elements can be further upstream or located in regions within the gene or downstream of the gene (Jeong et al., 2006; Laxa, 2016; Gallegos and Rose, 2019), (Weber et al., 2016). Models that attempt to predict changes in transcript abundance in response to stress likely should utilize motifs that may predict transcriptional changes as well as motifs within the RNA that influence stability of transcripts. We used a variety of different parameters to identify or filter the sequences that were used to discover enriched motifs or predict expression responses. We noted several important take-away messages from our attempts to document enriched motifs or predict responses to stress in B73. The use of larger regions and the inclusion of sequences within the transcribed region resulted in the discovery of more motifs, which is expected. More importantly, we also found that using these larger regions also improved the prediction accuracy of the models. However, we find that just using more sequence does not necessarily improve the prediction accuracies. Studies have highlighted the value of using chromatin modifications to reveal the location of important regulatory regions, especially in species with large complex genomes (Oka et al., 2017; Ricci et al., 2019; Crisp et al., 2020). Predictive models that only use the presence of motifs within unmethylated regions or accessible regions provide improvement in performance for some groups of genes even though this removes >50% of the sequence that is methylated. In contrast, the use of regions defined as accessible chromatin in a previous study (Ricci et al., 2019) did not improve prediction accuracies. Recent work has suggested that unmethylated regions may provide a catalogue of potential regulatory elements in plants (Crisp et al., 2020). Many regions that contain stress-specific regulatory elements are likely unmethylated even in control conditions while many of the regions of accessible chromatin may be more dynamic and only become accessible in specific conditions (Zeng et al., 2019; Wang et al., 2020a; Parvathaneni et al., 2020).

We had hypothesized that motif discovery and predictive models might be substantially improved by using time-course data to identify genes with different dynamics of stress response instead of simply using all up- or down-regulated genes. This was supported in most cases, but not in all. The predictions of genes that are down-regulated in response to cold were improved as well as the genes with transient up-regulation in response to heat. This may suggest that there are specific motifs that confer these patterns, and that by grouping these genes with other patterns of expression reduces the ability to predict the responses. In contrast, the performance for down-regulation in response to heat is not improved by focusing on specific patterns of response. Generating time-course data generally requires substantially more resources and our data for cold/heat responses suggest limited overall benefits in developing predictive models based on this data.

### Predictions of response to the environment across genotypes

A key goal in this work was to investigate the variation in response to abiotic stress in different maize genotypes and to assess approaches for predicting this variation. Local adaptation of plant populations likely involves changes to the *cis*-regulatory elements that allow for gene expression responses to environmental challenges. There are open questions about the nature of molecular variants that will create *cis*-regulatory changes in responsiveness to stress. SNPs within critical motifs could change the response but this is more likely to result in loss of a response rather than a novel responsiveness. Alternatively, structural variants including deletions or transposon insertions could provide novel elements or change the spacing between potential *cis*-regulatory elements and the TSS. We identified hundreds of genes that exhibit response to abiotic stress in one genotype but not another. These likely include variable responses that are due to both *cis-* and *trans*-regulatory variation, which are both of interest. The *trans*-regulatory variation for response to stress could indicate changes in specific TFs or signaling pathways among genotypes. Our study was able to identify examples of trans-regulatory variation but did not provide the ability to map the source of these variations. The *cis*-regulatory variation for response to stress indicates that one allele contains the elements necessary for response while the other does not. Further studies of these examples could provide insights into the molecular basis for changes in response to the environment.

The availability of genes with documented variation for response to heat or cold stress provided an opportunity to assess how well we could predict variation within germplasm for these responses. Ideally, we would be able to use data from a single genotype to effectively predict expression responses in other genotypes as the value of predictive models is partially due to the reduction in what data needs to be collected. However, we found substantially lower prediction accuracies when models trained solely on B73 expression responses were used to predict Mo17 or W22. If we use expression responses from all three genotypes, we achieve much higher prediction accuracies. However, these models are likely over-fitted as many genes show similar responses across the three genotypes and sequence similarity of these alleles will inflate accuracies. A truer estimate of the accuracy for cross-genotype predictions arises from focusing on the ability to accurately predict genes with variable response. While we were able to predict some of these examples of variable response the rates are relatively low (~20%). Most genes that have variable expression responses between genotypes were predicted to respond in both genotypes. We also attempted to generate models that were trained only using genes that have a variable response to stress in which the negative control set reflected the alleles that did not respond to stress. These models were not able to predict any better than a random guess suggesting difficulties in truly training to identify variable responses. Substantial improvements in cross-genotype prediction accuracies for genes with a variable response would be needed before this could be valuable in generating information that could be used to inform breeding decisions.

## Acknowledgments

We thank Peter Hermanson for assistance with plant growth, sampling and library preparation. We thank the Minnesota Supercomputing Institute at the University of Minnesota (http://www.msi.umn.edu) for providing resources that contributed to the research results reported within this article. This study was funded by the National Science Foundation (grants IOS-IOS-1444456, IOS-1546899 and IOS-1733633).

## Author Contributions

PZ, TAE, PAC, KG and NMS designed the research and generated the biological materials. PZ, ZAM and EM analyzed data. FG-C, ZL and EG contributed analytic tools. PZ and NMS wrote the paper.

## Methods

### Plant material and experimental design

Three experiments were carried out to study maize seedlings’ response to cold- and heat-stress treatments (Figure S1). Experiment 1 (TC) is a time-course experiment looking at cold- and heat- response at 9 different time points (0, 0.5, 1, 1.5, 2, 3, 4, 8 and 25 hours) after stress treatment for three maize inbreds (B73, Mo17 and W22). Experiment 2 (HY) sampled three inbred parents (B73, Mo17 and W22) as well as their hybrids (B73xMo17, B73xW22, Mo17xW22) each with three biological replicates but only at three time points under control, cold and heat conditions. Experiment 3 (NM) assayed a diverse genotype panel of 25 maize inbreds at two time points under control and cold conditions. Samples at time zero were re-used for control, cold and heat conditions in Experiment 1 and 2. Due to a low germination rate of W22 seedlings in Experiment 1, three time points (1.5, 3 and 8 hours after treatment) were skipped for W22 samples under control, cold and heat conditions, leading to a total of 66 samples in Experiment 1.

The three experiments were performed from late May to early July in 2019. In each experiment maize seeds of selected genotypes were imbibed for 24 hours in distilled water and grown in growth chambers at 30℃/20℃ under 16h/8h light-dark cycles to V2/V3 stage (day 9) for stress treatments and collection of the V2 leaves. For all three experiments we started stress treatment (cold or heat) at 2 hours after dawn on day 9 (i.e., time zero), and took samples from the control, cold and heat groups simultaneously at described time points. The temperature settings for control, cold and heat conditions were 30℃/20℃, 6℃/2℃ and 39℃/29℃, respectively. For each replicate V2 leaves from three maize seedlings were collected and pooled. Experiment 1 (TC) and 3 (NM) only have a single replicate while Experiment 2 (HY) has three replicates. A total of 292 samples were collected for profiling: 66 samples in experiment 1 (TC), 126 samples in experiment 2 (HC) and 100 samples in experiment 3 (NM). One sample did not meet the minimum cDNA content requirement during RNA-Seq library preparation and was discarded (Table S1).

### RNA-Seq data processing

Sequence libraries were prepared using the standard TruSeq Stranded mRNA library protocol and sequenced on NovaSeq 150bp paired end S4 flow cell to produce at least 20 million reads for each sample (Table S1). Both library construction and sequencing were done in the University of Minnesota Genomics Center. Sequencing reads were then processed through the nf-core RNA-Seq pipeline (Ewels et al., 2020) for initial QC and raw read counting. In short, reads were trimmed using Trim Galore! and aligned to the B73 maize reference genome (AGPv4, Ensembl Plant release 32) using the variant-aware aligner Hisat2 (Kim et al., 2015) and a graph index incorporating 90 million common maize variants to account for mapping bias. Uniquely aligned reads were then counted per feature by featureCounts (Liao et al., 2014). Raw read counts were then normalized by library size and corrected for library composition bias using the TMM normalization approach (Robinson and Oshlack, 2010) to give CPMs (Counts Per Million reads) for each gene in each sample, allowing direct comparison across samples. CPM values were then normalized by gene CDS lengths to give FPKM (Fragments Per Kilobase of exon per Million reads) values. Hierarchical clustering, principal component analysis and t-SNE clustering were used to explore sample cluster patterns and to remove or substitute bad replicates. Two samples were found to have genotype labels switched and were thus corrected (TC19: Mo17 and TC20: B73). The two missing time points in W22 (3hr and 8hr) were imputed from the two neighboring time points using linear imputation. Two HY samples at 25h time points (HY91, HY93) were also removed due to poor correlation with other biological replicates and substituted by two TC samples (TC64, TC66) and two NM samples (NM76, NM100) with matching time points and genotype.

### Identification of differentially expressed genes and characterization of genotypic difference

We used the replicated HY experiment to call differentially expressed genes (DEGs) between stress-treated samples and control samples. Since each time point in the experiment has two potential controls: one time 0 control and one control at the matching time point, two comparisons were made: one comparing the stress-treated samples at 1 or 25 hours with control samples at time zero (0 hour), and one comparing the treated samples with control samples at matching time point (1 or 25 hours). This led to two sets of DEGs identified for each genotype at each time point, the overlap between which were considered true stress-responsive genes and kept for downstream analysis. All statistical tests were done using the DESeq2 package in R (FDR adjusted P-value < 0.05 and a minimum fold change of 2) (Love et al., 2014).

To characterize the genotypic effect in each gene’s response to cold and heat stress, we made use of the Generalized Linear Model fitting framework in DESeq2 (Love et al., 2014). An interaction term between treatment condition and genotype was introduced and the model design was formulated as *Design = ~ Genotype + Condition + Genotype:Condition*. Using B73 as a baseline, we identified genes showing significantly different responses in Mo17 and W22 (compared to the response of B73) as well as those showing different responses between Mo17 and W22. In each comparison, significance is defined as FDR adjusted P-value < 0.05 and a minimum fold change (between two genotypes’ response to cold/heat treatment) of 2. Similar to calling DEGs for each genotype, this between-genotype test was also done twice - one using time 0 control samples and one using control samples at matched time points. A gene needs to show significance in both comparisons in order to be claimed as having a significant genotype effect.

The list of genes showing genotypic effect in each of the three pairwise comparisons (B73 to Mo17, B73 to W22 and Mo17 to W22) in each condition (cold_1h, cold_25h, heat_1h and heat_25h) were further grouped into 27 categories based on their response status in the three genotypes. We focused on the categories where stress response (either activation of repression) was lost in one or two genotypes, and used these as candidates to validate the cis-regulatory motifs we discovered.

### Identification of cold- and heat-responsive co-expression clusters

We used the time-course experiment to identify co-expression clusters following cold or heat stress. Our first analysis only looked at genes showing differential expression in the HY experiment at one of the two time points in B73. Since each time point of the time-course experiment has one sample in the treatment group and one control sample at matching time point, we explored three ways to construct the raw expression matrix for clustering: 1) CPM values of the treated sample at each time point; 2) CPM differences between treatment and control; and 3) log2 ratio of CPMs between treatment and control. We found that option 2 generally led to both interpretable and an ideal number of clusters. Expression levels for the three missing time points in W22 were imputed linearly using two neighboring time points. Distance-based hierarchical clustering was then used to discover gene co-expression clusters. The matrix of gene expression differences (CPM_stress_ - CPM_control_) is normalized using variance stabilizing transformation (VST) (Anders and Huber, 2010). The Pearson correlation coefficient-based distance matrix is then obtained and used for hierarchical clustering (method = “ward.D2”). The resulting gene tree was cut using the “cutreeDynamic” function (deepSplit = 3, minGap = 0) to yield 10-30 clusters along with their eigengenes (i.e., first principal component of the standardized expression vectors). Clusters with very similar eigengenes were then merged using different parameters (cutHeight = 0.1/0.15/0.2/0.25/0.3), results of which were visually inspected to determine the best cutting height. A list of 23 and 18 clusters containing 206-2307 genes were ultimately obtained for cold and heat responsive genes (Figure S3), from which 4 and 3 clusters were picked for downstream analysis (Figure 2A).

### Identification of enriched motifs in upstream/downstream regions of stress-responsive gene clusters

Motif mining was performed using different sets of genes including the list of all up- and down- regulated genes under cold or heat stress as well as specific co-expression clusters showing distinct time-course expression patterns (e.g., early or late up-regulation). STREME was run on each gene set to identify enriched motifs (8-20 bp ungapped k-mers) using genes not showing differential expression (P.adj > 0.05, less than 1.5 fold change in average expression) as negative controls (Bailey, 2011). For each gene set we explored the effect of different search spaces including different promoter size (0.5kb, 1kb and 2kb) around the transcription start site (TSS), or a combination of these regions) as well as a methylation- or accessibility-masked promoter space (i.e., only retain regions classified as unmethylated (Crisp et al., 2020) or accessible in leaf (acrL) or ear (acrE) (based on (Ricci et al., 2019)) (Figure 3A).

Known TF binding motifs from arabidopsis and maize were obtained from multiple sources (Weirauch et al., 2014; Ricci et al., 2019; Tu et al., 2020). Position weight matrices were either directly downloaded from the cis-BP website or built from called DAP-Seq peaks using GEM (Guo et al., 2018b). All motifs identified by STREME were compared to these public TF PWMs by calculating an all-against-all pairwise similarity matrix using bioconductor package universalmotif (method=“PCC”, min.mean.ic=.0, min.overlap=5, score.strat=“a.mean”) (Tremblay). Two rounds of hierarchical clustering (method=‘average’) were then performed to cluster motifs into motif groups. Any STREME motifs grouping with a public TF motif were assigned a “known” status and the corresponding TF label. Motif groups containing only STREME-identified motifs and no public TF motifs were assigned a “novel” status.

#### Train machine learning models to predict stress response based on cis-regulatory elements

Different sets of genes (e.g., all up- and down- regulated genes under cold or heat stress, early or late up-regulation under cold or heat stress, etc) were selected as positives and non-DE genes were selected as negatives to train machine learning models to predict stress responsiveness. Several machine learning algorithms (SVM, KNN, logistic regression, gradient boost, xgboost, random forest) were evaluated but random forest was selected since it consistently gave the best performance (data not shown). Downsampling was done prior to training to achieve balance between label groups. To train each model we first split data into 80% training set and 20% test set. Training was done within the 80% training set using 10 fold cross validation and evaluated using the 20% test set for performance. Model hyperparameters (number of predictors, number of trees, minimum number of data points in a node, etc.) were determined using a grid search algorithm (“tune_grid” function) implemented in R package “tidymodels”. Training for each co-expression cluster was repeated 100 times and the mean, median and standard deviation of performance metrics (F1 score, AUROC, AUPRC) were reported. After assessing the effect (F1 score) of different model training parameters (Figure 4, Figure S9) we determined a set of optimal parameters: “+/−2k TSS & +/−2k TTS”, “UMR”, “top100” features and using motif copy number as feature representation.

#### Characterization of genes showing stress-responsive cis- or trans-regulatory patterns

To classify gene expression levels into different regulatory categories, for each gene, we introduce the following notation (using B73 and Mo17 for example):

a_i_ = expression of the gene in the i^th^ B73 parent under control condition
b_i_ = expression of the gene in the i^th^ Mo17 parent under control condition
c_j_ = number of reads mapping to the B73 allele in the j^th^ F1 hybrid under control condition
d_j_ = number of reads mapping to the Mo17 allele in the j^th^ F1 hybrid under control condition
a_i_’ = expression of the gene in the i^th^ B73 parent under stress condition
b_i_’ = expression of the gene in the i^th^ Mo17 parent under stress condition
c_j_’ = number of reads mapping to the B73 allele in the j^th^ F1 hybrid under stress condition
d_j_’ = number of reads mapping to the Mo17 allele in the j^th^ F1 hybrid under stress condition

Here i and j take values between 1 and 4. Subsequently, we make the following distributional assumptions:

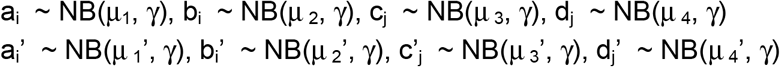

The dispersion parameter (℃) for each gene was estimated using the “estimateDispersions” function within DESeq2 with *fitType = ‘parametric’* option (Love et al., 2014). The marginal distributions of a_i_ are negative binomial. Subsequently, different constraints upon the parameters can be imposed to describe the following biological situations:

First define: *d* μ _1_ = μ _1_’ - μ _1_, *d* μ _2_ = μ _2_’ - μ _2_, *d* μ _3_ = μ _3_’ - μ _3_, *d* μ _4_ = μ _4_’ - μ _4_
conserved: *d* μ _1_ = *d* μ _2_ and *d* μ _3_ = *d* μ _4_,
unexpected: *d* μ _1_ = *d* μ _2_ and *d* μ _3_ ≠ *d* μ _4_,
cis: *d* μ _1_ ≠ *d* μ _2_ and *d* μ _1_/*d* μ _2_ = *d* μ _3_/*d* μ _4_,
trans: *d* μ _1_ ≠ *d* μ _2_ and *d* μ _3_ = *d* μ _4_,
cis & trans: *d* μ _1_ ≠ *d* μ _2_ and *d* μ _1_/*d* μ _2_ ≠ *d* μ _3_/*d* μ _4_

Each gene was allocated into one of these four categories by first fitting the four models described above to the data by maximizing the likelihood function, then calculating the Bayesian Information Criterion (BIC) to determine which of the four models best fitted the data for each gene.

#### Identification of putative variants regulating cold-responsive expression in a panel of 25 maize genotypes

Starting from a set of ~2000 genes showing variable response patterns at 25 hour after cold treatment between B73, Mo17 and W22, we checked their response patterns in a wider set of 25 maize genotypes. The levels of activation / repression (log2 fold change) for different maize inbreds were plotted. Bi-modal distribution of log2 fold changes were searched with one cluster centering around 0 suggesting no DE while the other cluster departing from 0 suggesting significant DE. A total of ~500 genes were identified showing significant bi-modal distribution of their log2fc values among the 25 NAM genotypes that contain at least one non-DE genotype and a DE genotype. Genomic sequences of the 25 maize genotypes were then aligned to B73 and bi-allelic variants within 2kb of each gene were extracted to perform a local association test with the cold response pattern (i.e., treating the bi-modal distribution as one group of stress-responsive genotypes and another group of non-stress-responsive genotypes). We used PLINK to perform a standard chi-square test for each gene with multiple testing correction (“--assoc -- adjust”). Significantly associations were determined by requiring the Bonferroni corrected P-value < 0.05. Since many variants are in high linkage disequilibrium, for visualization purpose we identified haplotype blocks using the default procedure implemented in PLINK (“--blocks-min-maf 0 --blocks-strong-highci 1 --blocks-inform-frac 0.8”).

## Data Availability

Raw RNA-Seq reads have been deposited in NCBI Sequence Read Archive (SRA) under accession PRJNA482146. All source codes used for quantification, normalization, statistical testing and machine learning training and evaluation are available on Github: https://github.com/orionzhou/stress. The processed data sets including gene lists of each stress responsive pattern, lists of genes under cis/trans stress responsive regulation, lists of enriched motifs found in each co-expression cluster are deposited at http://hdl.handle.net/11299/212030

## Supplemental figure legends

Figure S1. Experimental design used for generation of RNA-Seq data. (A) Temperature readings from a sensor that was with the plants throughout the experiment and shows the time of light / dark (gray shading indicates time without lights). (B-D) Three different experiments were conducted to assess the changes in gene expression in response to cold or heat stress in 14 day maize seedlings. In experiment 1 (time course - TC, panel B) one replicate for three inbreds was collected at each of the indicated time-points. In experiment 2 (inbred - hybrid - HY, panel C) three biological replicates were sampled from three maize inbreds and their F1 hybrids at time 0 and two time points during the stress. In experiment 3 (panel D) a single biological replicate was sampled for a panel of 25 genotypes under control and cold conditions at two time points.

Figure S2. Hierarchical (A) and t-SNE (B) clustering of all samples from the time course (TC) experiment under control, cold and heat conditions. (A) The samples showed strong clustering by genotype (indicated by color of labels) and within each genotype the samples tend to cluster by treatment. (B) t-SNE plot shows similar clustering based on both genotype and treatment.

Figure S3. Expression profiles of cold- and heat-responsive gene clusters. B73, Mo17 and W22 genes that exhibit significant DE after 1h or 25h stress treatment were used to perform co-expression clustering based on their time-course expression pattern (see methods). The median expression level of control and stress conditions for the genes within each module is shown and the number of B73, Mo17 and W22 genes in each module is indicated. Ranges at each time point represent 25%-75% quantile expression levels.

Figure S4. Transcription Factor Family Enrichment in Select DEG Modules. The TFs present in the overall set of up-regulated genes for cold (A) or heat (B) as well as TFs that are present in specific modules were assessed for potential enrichment of specific TF families. Numbers in parentheses indicate total group size, while numbers in tiles indicate how many DEGs of specific TF families were identified in each module. For modules including at least 3 TF members in a TF family, enrichment was assessed through hypergeometric test (p < 0.05). Colors in tiles indicate level of enrichment (-log10(p.value)). Transcription factor family assignments were pulled from Grassius TFDB (Yilmaz et al., 2009).

Figure S5. Characterization of maize Heat Shock Factors (HSF) response to heat stress. Time course expression pattern in B73 was shown for all 29 maize HSFs including: (A) 5 HSFs exhibit very strong activation at 0.5h of heat stress, but return to relatively normal levels by the 1h time point. (B) Four HSFs classified as transient activation; (C) One HSF was significantly differentially expressed, but not assigned to a specific cold-upregulated module. (D) Ten HSFs are expressed but not assigned into a cold up-regulation module. Genes were considered expressed if expression was ≥ 5 cpm in least one treatment x time point.

Figure S6. Expression profile of TFs that have previously been reported to play a role in maize response to cold. Out of 235 previously reported cold stress-related TFs, 109 TFs were expressed with ≥ 5 CPM in B73 in at least one sample (treatment x time point), while 43 were differentially expressed in response to cold stress and assigned into cold up-regulated modules including: (A) late activation, (B) early activation, (C) differentially expressed (DE). DE cold up-regulated genes were identified in the hybrid experiment (3 replicates, 1h and 25h after stress) and TF gene response to cold stress in the time-series (1 replicate, 9 time points) may not depict cold up-regulation for each DE gene.

Figure S7. Number of enriched motifs found using B73 promoter spaces (“B model”) or B73/Mo17/W22 promoter spaces (“BMW model”) of different sizes and epigenetic filters. Co-expression modules as well as the entire set of cold-/heat- up-/down-regulated genes were searched for enriched motifs with varying lengths of promoter space and different filters based on methylation or chromatin accessibility. Identified motifs were compared with known TF binding motifs (cis-BP) and clustered based on sequence similarity. Numbers in each cell represent the number of non-redundant motifs found using the full set of sequences (raw), using only the unmethylated regions (umr) or using only accessible chromatin regions in leaf (acrL) or ear (acrE) inside different sized promoter spaces. Color indicates the number of motifs identified (the darker, the more motifs found) and corresponds to the number in each cell. Numbers in parentheses on the left side of plot indicate the number of genes in each co-expression module used for motif mining, while numbers on the right side of plot indicate the total number of non-redundant motifs found for each set of genes after sequence clustering.

Figure S8. Enriched motifs include known transcription factor binding sites (TFBSs) as well as novel motifs. For each set of DEGs up to the top 40 most enriched motifs found using B73 promoter space (“B model”) or B73/Mo17/W22 promoter sequences (“BMW model”) are shown (p-value for enrichment is indicated by color). Some sets of DEGs have less than 40 motifs and only the significant ones are shown. If the enriched motif matched a previously characterized TFBS (Pearson correlation coefficient > 0.8) the name of the transcription factor is shown. In each case, there are a mixture of previously characterized motifs and novel motifs. TTS motifs with a “*” sign represent motifs are not present in the top40 TSS motifs identified in the same list of genes.

Figure S9. Meta plots of selected stress-responsive motifs in gene promoters. Two to four motifs from each of the seven co-expression clusters (cold early activation, cold late activation, etc.) with their relative locations in the TSS:−/+2k region cut into 40 bins. The occurrence of each motif in these bins were recorded and the proportion of genes containing the motif in each bin is plotted.

Figure S10. Performance (Area Under ROC Curve, AUROC) of machine learning models predicting cold and heat responsive expression. Models were trained to predict expression responses to heat or cold using: (A) different sizes of promoter sequences surrounding the TSS; (B) using “binary encoding” (0/1) or “quantitative encoding” (0/1/2/…) of motifs and and (C) using different numbers of mostly enriched motifs as input features. The number of non-redundant motifs used for model training are indicated on top of each set of bars in (C). In each training the average AUROC (N=100 model runs) is show along with the standard deviation.

Figure S11. Relationship between motif enrichment level and feature importance score in different sets of stress responsive genes. Motif enrichment levels are determined by a hypergeometric test using motif occurrences in positive and negative gene sets and ranked from most significant to least significant (x-axis). Feature importance scores reported by 100 permutations of each random forest model training are shown on the y-axis with error bar indicating 25-75% quantiles. The top 5 feature importance scores are labelled with motif names (known motif) or consensus sequences (novel motif). All feature importance score estimates are based on the models trained using “+/−2k”, “UMR”, “top200”, “binary” parameters.

Figure S12. Comparison of heat- and cold-response gene expression in B73, Mo17 and W22. (A) At each time point for cold or heat stress we identified DEGs in all three genotypes and show the overlap of DEGs. The actual expression levels in all three genotypes for the non-redundant set of up-regulated genes is shown in (B). The genes are classified based on response in B73, Mo17 and W22 (0 indicates not DE and 1 indicates DE in the three genotypes such that ‘111’ indicates DE in B73, Mo17 and W22 while ‘010’ would indicate DE in Mo17 but not B73 or W22. Note that for some genes classified as having variable responses there are actually similar changes in all genotypes but there are other examples with clear response on some genotypes but not in others.

Figure S13. Characterization of genes with variable stress responsive patterns among inbreds. For each pair of genotypes we identified genes that show significantly stronger (or weaker) response to cold/heat at 1h or 25h time points using a generalized linear model (see methods). (A) Scatterplot showing the log2 fold-change (stress / control at a time point) for compared inbreds (A and B) in x- and y-axis. Above the diagonal line are genes showing stronger cold response in genotype A while those below the diagonal line represent genes showing stronger genotype B response. Different colors indicate groups of genes with different DE status (e.g., red genes are ones up-regulated in both inbreds (but showing significantly different response), green are genes only up-regulated in genotype B). (B) The number of genes in each category shown in (A). For each class the response in the two genotypes (A and B) is indicated as up-regulated (“+”), down-regulated (“-”) or not DE (=).

Figure S14. *Cis/trans* characterization of genes showing different stress response among inbreds. For the subset of genes classified as showing significantly different responses among inbreds that also had SNPs we assessed allele-specific expression in the F1 hybrid. (A) For each pairwise genotype comparison the proportion of allele 1 change in stress vs control of the F1 (x-axis) was compared to the proportion of the change in expression in the parental genotypes (y-axis). A maximum likelihood model was applied to classify *cis-* and *trans-* inheriance patterns and shown in different colors. (B) The number of genes classified into each type of regulatory pattern in each pairwise comparison.

Figure S15. Identification of *cis*-regulatory variants associated with variable cold responsive pattern in a panel of 25 maize genotypes. (A) Consistency of log2 fold changes for each gene called between two experiments: the replicated HY experiment with 3 maize inbreds (x-axis) and the NM experiment with 25 maize genotypes (y-axis). Point density is shown to avoid overplotting. Linear regression is performed and the adjusted R2 values are marked. (B) *Cis*-regulated variable response genes are enriched in having significantly associated local variants. Shown are proportions of genes with detected local associations for variable response genes under different regulatory patterns.

Figure S16. *Cis*-regulatory variation associated with cold responsive patterns in a respiratory burst oxidase (rboh4, panel A) and a dihydroflavonoid reductase (dfr1, panel B). Heatmap shows bi-allelic variants (SNP and short indels) within 2kb of the gene with yellow indicating reference (B73) allele and blue indicating alternate allele. Log2 fold change of each genotype is shown on the right of the heatmap. Below the gene structure plot are locations of the most (top30) enriched cold-responsive motifs (red dots) with the motif name marked, as well as haplotype blocks (dark blue segments) identified using PLINK. Boxplot on the right shows the top associating variant and the log2fc distributions of genotypes carrying the two alleles.

